# Transformers enable accurate prediction of acute and chronic chemical toxicity in aquatic organisms

**DOI:** 10.1101/2023.04.17.537138

**Authors:** Mikael Gustavsson, Styrbjörn Käll, Patrik Svedberg, Juan S. Inda-Diaz, Sverker Molander, Jessica Coria, Thomas Backhaus, Erik Kristiansson

## Abstract

Environmental safety assessments, as mandated by many regulations, require that toxicity data is generated for up to three trophic levels, algae, aquatic invertebrates, and fish. Conducting these tests *in vivo* is resource-intensive, time-consuming, and causes undue suffering. Computational methods are fast and cost-efficient alternatives, however, their adaptation in regulatory settings has been slow, both due to low accuracy and narrow applicability domains. Here we present a new method for predicting chemical toxicity based on molecular structure. The method is based on a transformer, capturing structural features associated with toxicity, followed by a deep neural network that predicts the corresponding effect concentrations. After training on data from tens of thousands of exposure experiments, the model shows high predictive performance for each of the three trophic levels. Compared to commonly used QSAR methods, the model has both a larger applicability domain and a considerably lower error. In addition, training the model on data that combines multiple types of effect concentrations further improves the performance. We conclude that transformer-based models have the potential to significantly advance computational predictions of chemical toxicity and make *in silico* approaches a more attractive alternative when compared to animal-based exposure experiments.

## 1 Introduction

Chemical pollution is a major contributor to the declining ecological status of European surface waters (Malaj et al., 2014; Posthuma et al., 2019; van de Meent et al., 2020). It also constitutes a planetary boundary that cannot be transgressed without risking large irreversible environmental changes (Diamond et al., 2015; Persson et al., 2022; Rockström et al., 2009). Several adverse environmental effects have been directly associated with chemical pollution, such as the extreme decline in vultures in India (Oaks et al., 2004; Swan et al., 2006) and the general decline in bee populations in the Western world (Sgolastra et al., 2020). Chemical pollution also negatively affects humans, with an estimated cost of disease of 157 billion € and 340 billion $US for the EU and US, respectively (Attina et al., 2016; Trasande et al., 2015). These adverse effects are, primarily, caused by chemicals that due to a lack of accurate data were originally considered safe, but once released in sufficient amounts caused damage to both human and environmental health.

To ensure that chemicals are used in a safe and sustainable way, increasingly stringent legal systems have been implemented over time (Botos et al., 2019; van Dijk et al., 2021). Environmentally safe concentrations are determined based on whole organism exposures to chemicals. For the aquatic compartment, up to three trophic levels are considered; primary producers, primary consumers, and secondary consumers (represented by algae, aquatic invertebrates, and fish) (ECHA, 2008; EFSA, 2013). Currently, more than 2 million animals are sacrificed annually for regulatory purposes (European Commission, 2020), a number that is expected to increase due to the continuous expansion of the number of chemicals used in society (CEFIC, 2022; Llanos et al., 2019).

Computational methods have been suggested as fast and cost-efficient alternatives to whole-animal exposure experiments (Cherkasov et al., 2014; Muratov et al., 2020). This includes, in particular, quantitative structure-activity relationship (QSAR) methods that use regression or other predictive models (e.g., machine learning-based) to associate differences in chemical structures with changes in toxicological properties. Effects of larger structural alterations are, however, notoriously hard to predict, and traditional QSAR models are, therefore, typically developed using data that is highly stratified according to chemical structure, toxicological effects, and species, as well as exposure scenarios and endpoints. Due to the narrow applicability domains, multiple QSAR models are often required to make predictions for more than one single chemical class. Alternatively, machine learning techniques have been suggested as an approach to integrate large volumes of heterogeneous data into a single, more general, model (Jeong and Choi, 2022). For example, deep learning has been used to predict various biological activities, such as toxicity, based on chemical structures (Ciallella et al., 2021; Goh et al., 2017; LeCun et al., 2015; Mayr et al., 2016). However, no method suggested so far has the necessary accuracy and a sufficiently large applicability domain to make it applicable to most regulatory purposes. Indeed, existing computational methods have only been able to replace a very small proportion of whole-animal tests, and new approaches are, thus, needed to ensure that chemical regulation remains in pace with the increasing number of chemicals introduced (Llanos et al., 2019; Wang et al., 2020).

Recently, transformers, a deep learning methodology originally developed for natural language processing (Devlin et al., 2018; Vaswani et al., 2017), have been shown to be highly efficient at capturing information from biological and chemical structures (Chithrananda et al., 2020; Jumper et al., 2021). Transformers use self-attention, a mechanism that infers complex dependencies directly from data, and uses it to emphasize the parts that are deemed especially informative. This makes it possible to identify the structural features that are most important for accurate prediction of chemical toxicity. The rapid accumulation of toxicity data, stemming both from European legislations with a ‘no data no market’ philosophy (ECHA 2022), and the continuously expanding ecotoxicological research community (Kristiansson et al., 2021), has resulted in data from exposure experiments of tens of thousands of chemicals becoming available. This paves the way for advanced deep-learning methods, such as transformers, to improve the computational predictions of chemical toxicity.

Here, we describe a transformer-based model that uses chemical structure, together with exposure parameters, to predict chemical toxicity. The model shows high predictive performance for aquatic organisms from three trophic levels: algae, aquatic invertebrates, and fish. Compared to three commonly used QSAR-based methods the proposed model had markedly improved accuracy. The transformer-based model is also able to make predictions for all chemical structures in the dataset, while many compounds fall outside of the other methods applicability domains. Finally, the model performance is further improved by combining multiple endpoints within a single model. We conclude that transformers, trained on rich chemical toxicity data, significantly advance the computational prediction of chemical toxicity, making it an increasingly attractive alternative to whole-animal toxicity experiments.

## 2 Results

In this study, we present a deep learning model for predicting chemical toxicity, based on molecular structure (Figure 1). The model uses a transformer encoder to derive a numerical representation of the chemical structure, which is used as input to a deep neural network that, together with information on the effect, measured endpoint, and duration of the exposure, predicts an associated effect concentration (EC_10_ and EC_50_) (Chithrananda et al., 2020; Vaswani et al., 2017). The transformer was trained on rich datasets of experimentally measured effect concentrations covering 2417 to 3492 unique chemical structures, for three organism groups (Table 1). Stratified sampling was used to reduce the influence of overrepresented chemicals and Bayesian optimization was used to estimate model hyperparameters (Falkner et al., 2018). Initially, two individual models were trained for each organism group, one for the prediction of EC_50_ and one for EC_10_ (S.I. Table 4, Methods). The code, data, and trained models are available via (https://github.com/StyrbjornKall/ecoCAIT) and (https://huggingface.co/StyrbjornKall).

**Figure 1:**
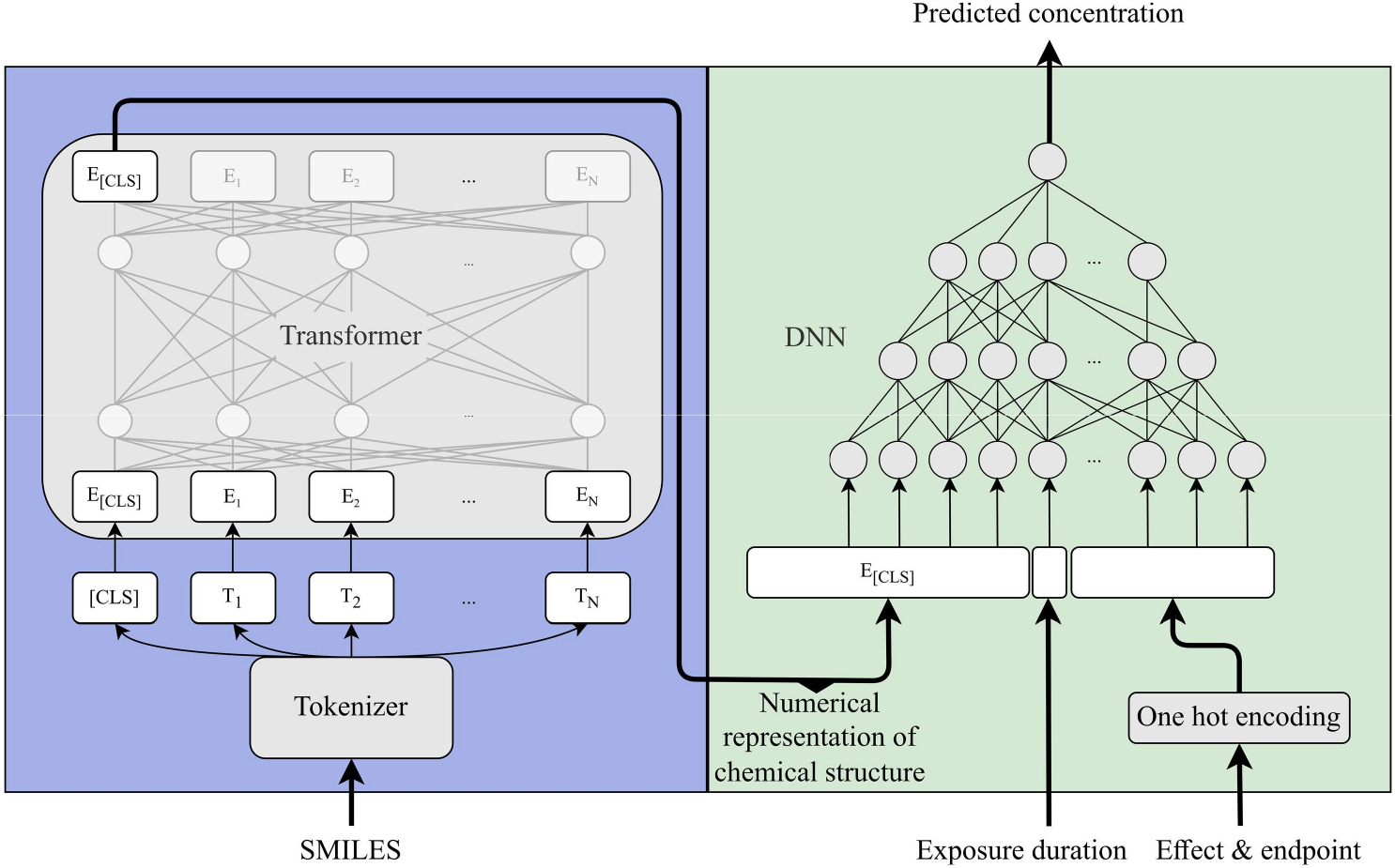
Model architecture. The model uses a pre-trained 6-encoder layer RoBERTa transformer (ChemBERTa) to interpret the SMILES into an embedding vector of dimension 768 (E_[CLS]_), representing the molecular structure with regard to its toxicity. The embedding vector is then amended with information on exposure duration, toxicological effect, and endpoint and used as input to a deep neural network. The network then predicts the associated toxicity in the form of an effect concentration (EC_50_ and EC_10_). The model parameters and hyperparameters were determined using Bayesian optimization (S.I. Table 4).

**Table 1:** Overview of the EC_50_ and EC_10_ datasets for fish, aquatic invertebrates, and algae. The datasets were used to train and validate the transformer-based model. The number of unique experimental setups is the number of unique combinations of chemicals, endpoints, effects, and exposure durations in each dataset.

| Dataset | Organism group | Endpoint | Effect | #Datapoints | #Unique SMILES | #Unique experimental setups | #SMILES responsible for over 50% of data | Concentration (mg/L) mean (95%,5%)* | Exposure duration (h) mean and SEM* |
| --- | --- | --- | --- | --- | --- | --- | --- | --- | --- |
| Fish EC<sub>50</sub> | Fish | EC<sub>50</sub> | MOR | 45017 | 2417 | 6733 | 59 | 23.8 (134, 0.004) | 86±0.7 |
| Fish EC<sub>10</sub> | Fish | EC<sub>00</sub>-EC<sub>10</sub>, NOEC | MOR, ITX, DVP, GRO, REP, MPH, POP | 23609 | 2951 | 8907 | 176 | 27.0 (168, 0.0002) | 555±9 |
| Aquatic invertebrates EC<sub>50</sub> | Aquatic invertebrates | EC<sub>50</sub> | MOR, ITX | 31653 | 3492 | 7958 | 67 | 23.5 (137, 0.0006) | 73±0.6 |
| Aquatic invertebrates EC<sub>10</sub> | Aquatic invertebrates | EC<sub>00</sub>-EC<sub>10</sub>, NOEC | MOR, ITX, DVP, REP, MPH, POP | 19170 | 2913 | 7817 | 117 | 20.1 (120, 0.0002) | 400±7 |
| Algae EC<sub>50</sub> | Algae | EC<sub>50</sub> | POP | 11909 | 2747 | 4113 | 181 | 27.6 (155, 0.007) | 96±1.2 |
| Algae EC<sub>10</sub> | Algae | EC<sub>00</sub>-EC<sub>10</sub>, NOEC | POP | 11384 | 2695 | 4153 | 170 | 17.7 (100, 0.002) | 234±7 |

\*Values were calculated from the log<sub>10</sub> transformed data used to train the models

We first examined how the numerical representation of the chemical structures corresponded to the chemical toxicity. Principal component analysis (PCA) of the 768-dimensional embedding vectors showed that the model organized the chemical structures in a continuous gradient for all models and all organism groups (Figure 2; S.I. Figure 2, S.I. Figure 3). For example, for fish, the EC_50_ model showed a clear ability to separate toxic and non-toxic chemicals (Figure 2a), demonstrating that the model properly captures toxicological information from the chemical structure. The EC_10_ models (Figure 2b, S.I Figure 2b, S.I. Figure 3b) followed similar trends but with more deviating chemical structures, reflecting a higher degree of uncertainty. It should be emphasized here that the numerical representation inferred by the transformer includes information about toxicity and closeness does thus not necessarily imply overall structural similarity.

**Figure 2:**
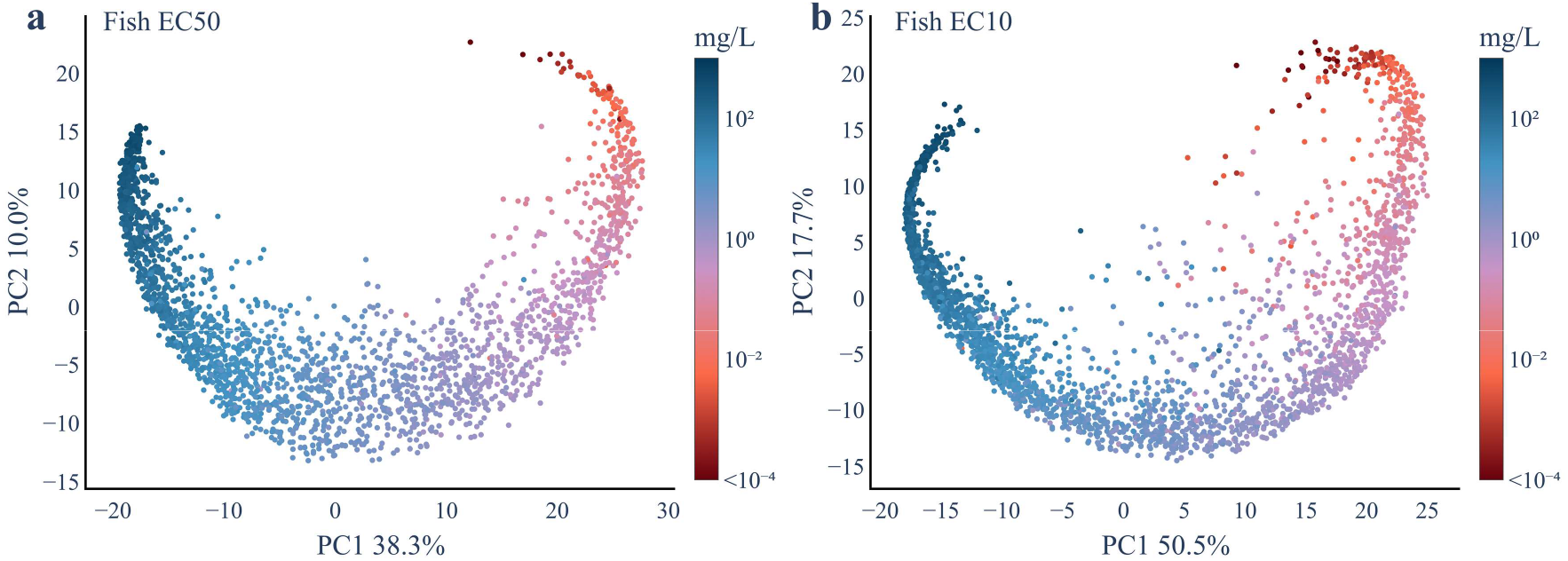
Principal component analysis (PCA) of the numerical presentation of chemical structure. The PCA shows a two-dimensional visualization of the structural representation inferred by the transformer from experimental exposure data on fish for a) EC_50_ (n = 2417) and b) EC_10_ (n = 2951). The PCA was calculated from the 768-dimensional embedding vector associated with each chemical structure. The color of each chemical structure is based on the median of the experimentally measured toxicity. The values at the x- and y-axis is the total percent of variation explained by the corresponding principal component.

**Figure 3:**
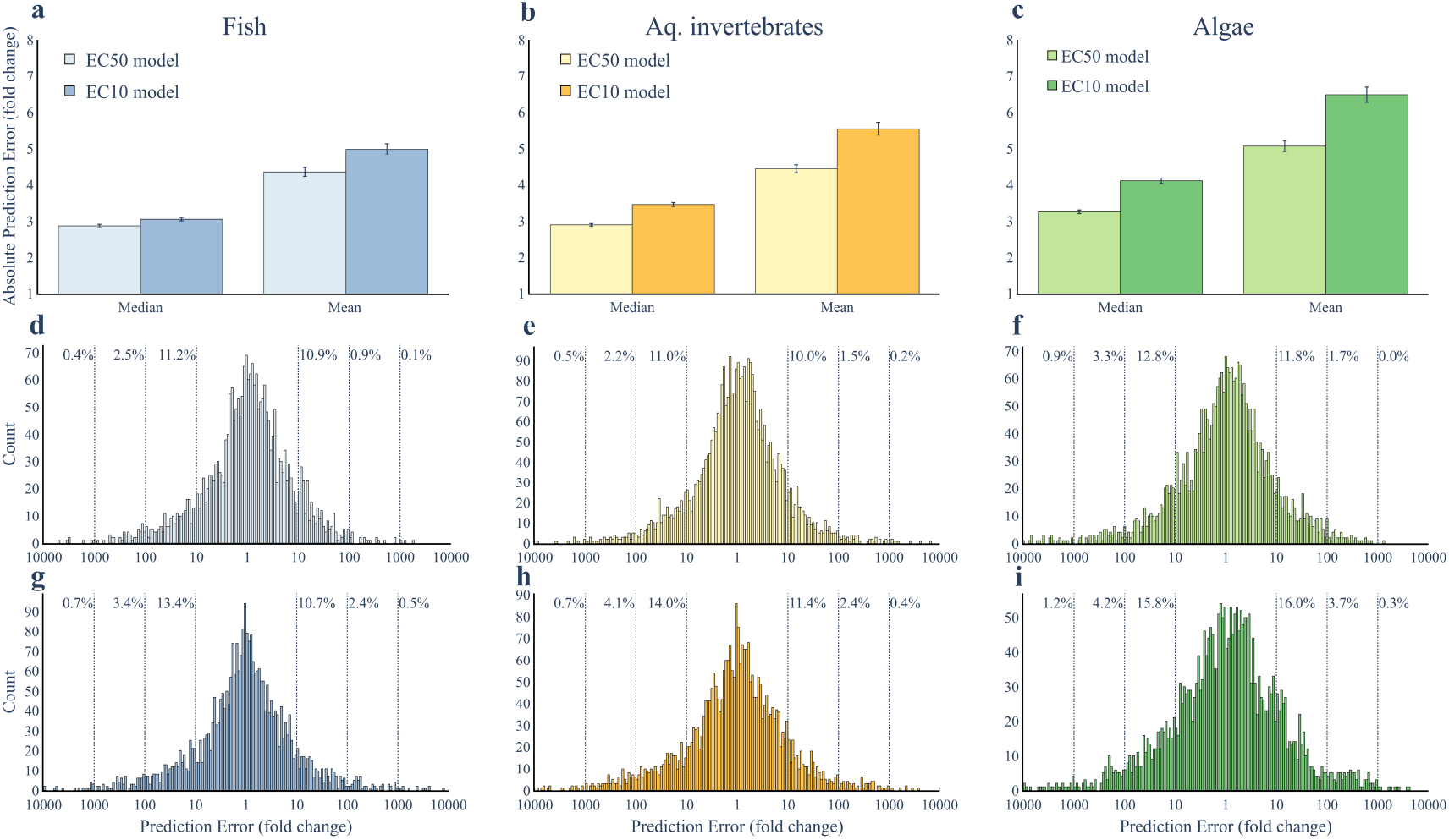
Model performance. Panels (a-c) show the performance as the median and mean absolute prediction error, measured as the absolute fold-change between predicted and experimental values, determined from the ten times repeated ten-fold cross-validations for the (a) fish EC_50_ model (n = 45017) and fish EC_10_ model (n = 23609), (b) aquatic invertebrate EC_50_ model (n = 31653) and aquatic invertebrate EC_10_ model (n = 19170), and (c) algae EC_50_ model (n = 11909) and algae EC_10_ model (n = 11384). The error bars show the median absolute deviation and the standard error of the mean for the respective prediction error. Panels (d-i) show histograms of residuals for the six models: (d) fish EC_50_, (e) aquatic invertebrates EC_50_, (f) algae EC_50_, (g) fish EC_10_, (h) aquatic invertebrates EC_10_, (i) algae EC_10_. The reported percentage values show the percentage of residuals that are larger than a factor of 10, 100, or 1,000, respectively.

The model performance was then evaluated using ten times repeated ten-fold cross-validation, where chemical structures were randomly divided into training and test datasets (Methods). The performance of the model was, thus, always measured using chemicals that were not included in the training and, hence, ‘new’ to the model. The highest predictive performances for EC_50_ were seen for fish and aquatic invertebrates, which showed, compared to experimental data, median error factors of 2.87 and 2.89, respectively (Figure 3a-b). The model performance for algae was slightly lower, with a median error factor of 3.25 (Figure 3c). A closer analysis of the residuals from the fish models showed that close to 80% of all predictions were within a factor of ten of the experimentally measured data and only 3.4% of the chemicals had prediction errors larger than a factor of 100 (Figure 3d). These numbers were similar for aquatic invertebrates (79.0% and 3.7%, respectively, Figure 3e) and Algae (75.4% and 5.0%, respectively, Figure 3f). Interestingly, the EC_10_ models were able to make predictions with almost the same accuracy (Figure 3a-c). The median errors were a factor of 3.05, 3.44, and 4.11 for the fish, aquatic invertebrates, and algae models, respectively. The corresponding proportion of chemicals deviating more than ten-fold were 75.9%, 74.6%, and 68.2% (Figure 3g-i). For all models, less than 1% of the chemical structures had a median error larger than a factor of 1000.

We then investigated if the performance would further increase if the model was extended to predict both EC_50_ and EC_10_ values. Three extended models were trained (S.I. Table 4), one for each organism group, and then evaluated using ten times repeated ten-fold cross-validation. Since we wanted to investigate if data for EC_50_ could improve the prediction of EC_10_, and vice versa, the model was allowed to be evaluated on chemical structure also present in the training data, as long as the predicted endpoint was different. Thus, the model was, e.g., allowed to train on a certain chemical and its corresponding measured EC_50_ values but only evaluated on the chemical’s EC_10_ data. The results showed that the model performance increased for all three organism groups. For the fish model, the decrease in the median error was 24.3% (2.17) and 21.6% (2.39) for EC_50_ and EC_10_, respectively (Figure 4). The increase in performance was even larger for aquatic invertebrates and algae, with decreases in error corresponding to 23.9% (2.20) and 39.5% (2.08) for aquatic invertebrates EC_50_ and EC_10_, respectively, and 38.5% (2.00) and 48.7% (2.11) for algae EC_50_ and EC_10_, respectively (S.I. Figure 4). As expected, the largest performance improvement was seen for chemicals where both EC_50_ and EC_10_ data was available. These chemicals had a median error of <2 for all organism groups and effect concentrations. This shows that the proposed model can accurately extrapolate between effect concentrations and, thus, use existing experimental data to further improve the predictive performance.

**Figure 4:**
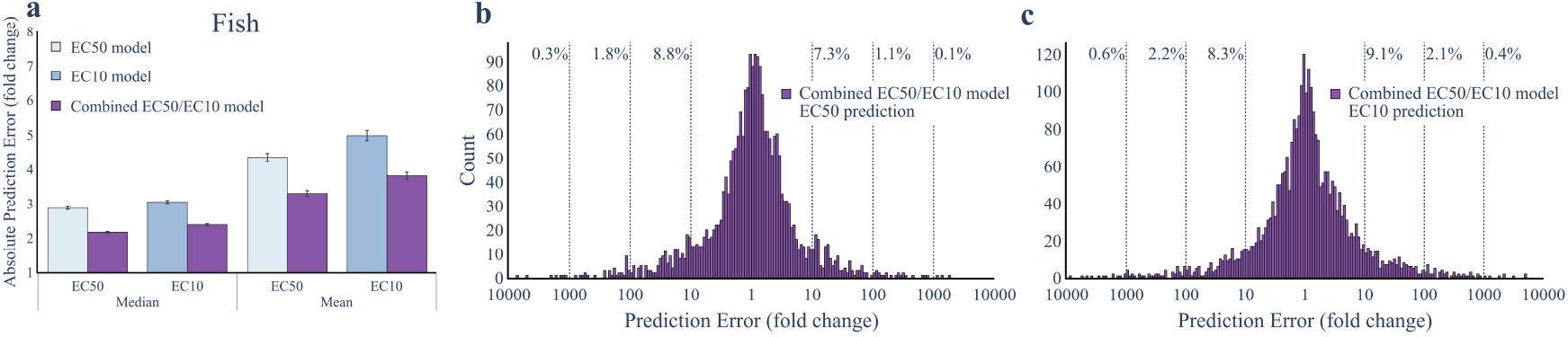
Combined model performance. Panel (a) shows the performance as the absolute median and mean prediction error, measured as the absolute fold-change between predicted and experimental values, determined from the ten times repeated ten-fold cross-validations for the fish EC_50_ model (n = 46104), fish EC_10_ model (n = 23609), and the fish model able to predict both EC_50_/EC_10_ (n = 69713). The error bars show the median absolute deviation and the standard error of the mean for the respective prediction error. Panels (b-c) show the histogram of residuals for the fish model able to predict both EC_50_/EC_10_ when predictions were evaluated on the EC_50_ and EC_10_ datasets. The reported percentage values show the percentage of chemicals that are erroneously predicted by a factor of more than 10, 100, or 1000.

The performance of the transformer-based model was then compared to three of the most commonly used traditional QSAR methods for the assessment of chemical toxicity in aquatic organisms (ECOSAR, VEGA, and T.E.S.T.) (Benfenati et al., 2013; Martin, 2020; Wright et al., 2022a). The transformer-based models were trained on a large and diverse set of chemicals and are, thus, not restricted to any specific chemical classes. Indeed, the transformer-based model could provide predictions for all structures included in the training datasets (n = 6508 unique structures). In contrast, a large proportion of these chemicals fell outside the applicability domains of the traditional QSAR methods (Figure 5). The largest differences in applicability domain were seen for EC_10_ (Figure 5a,c,d), where VEGA was only able to analyze 10% to 30% of the chemical structures (depending on organism group) while T.E.S.T. did not provide any predictions at all. Of the three QSAR methods, ECOSAR had the largest applicability domain for EC_10_, and it was able to predict toxicity for more than 75% of the chemical structures. Even when used in the most relaxed settings, under which ECOSAR and VEGA return predictions for chemicals outside of their applicability domains, these QSAR methods still failed to predict the toxicity of a large proportion of the chemicals (Figure 5b,d,f). Finally, there was a large overlap between the applicability domains of ECOSAR, VEGA, and T.E.S.T., likely reflecting that these models are developed and trained using partially similar data. This is reflected by the fact that the toxicity for between 15% and 24% of the chemicals could not be predicted by any of the traditional QSAR methods (S.I. Figure 5).

**Figure 5:**
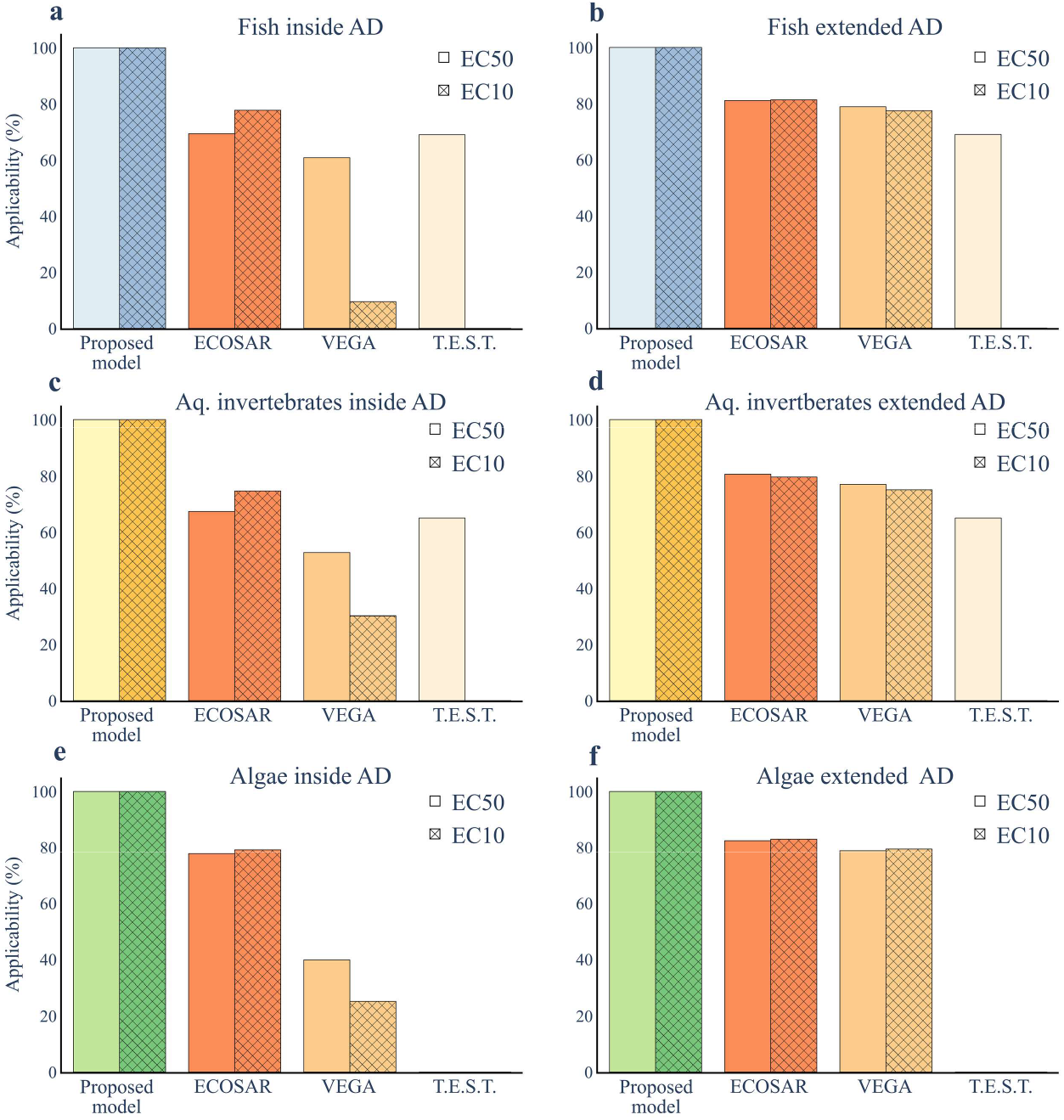
Comparison of applicability domains of evaluated models. The percentage of SMILES included in the six different datasets that ECOSAR, VEGA, and T.E.S.T. can predict toxicity values for compared to the applicability domain of the transformer-based models. The percentage is reported both for the chemicals that are within the reported applicability domain of the model (a, c. e) and chemicals that a prediction can be performed for, here called extended applicability domain (b, d, f). The comparisons are based on the total number of unique chemicals included in the current model (100%) for (a-b) fish EC_50_ (n = 2417), fish EC_10_ (n = 2591), (c-d) aquatic invertebrates EC_50_ (n = 3492), aquatic invertebrates EC_10_ (n = 2913), and (e-f) algae EC_50_ (n = 2747), algae EC_10_ (n = 2695).

The predictive performance of the transformer-based model was compared against the traditional QSAR methods (Figure 6). To make the comparison fair, we only included chemical structures that were inside the applicability domains of all methods and that were not included in any of the training datasets. For the predictions of EC_50_, the transformer-based model had the best accuracy for fish and aquatic invertebrates with a median error corresponding to a factor of 2.40 and 2.41, respectively, while ECOSAR had the best performance for algae with a median error of a factor of 1.41 (Figure 6a, Table 2). The difference between the methods was much larger for the EC_10_ predictions, where the transformer-based model had an error corresponding to a factor of 2.56, and 2.73, for fish and invertebrates, respectively. The second-best method, VEGA, had a predictive error four times as large, corresponding to a factor of 11.46 and 9.47 for fish and aquatic invertebrates, respectively (Figure 6b, Table 2). For algae, the differences were not as extreme, however, the transformer-based model had an error of a factor of 4.37, compared to 5.60 for ECOSAR, the second-best performing method. Additionally, the transformer-based model consistently had a lower error when evaluating the predictive performance individually for each included toxicological effect, (S.I. Figure 9, S.I. Figure 10).

**Figure 6:**
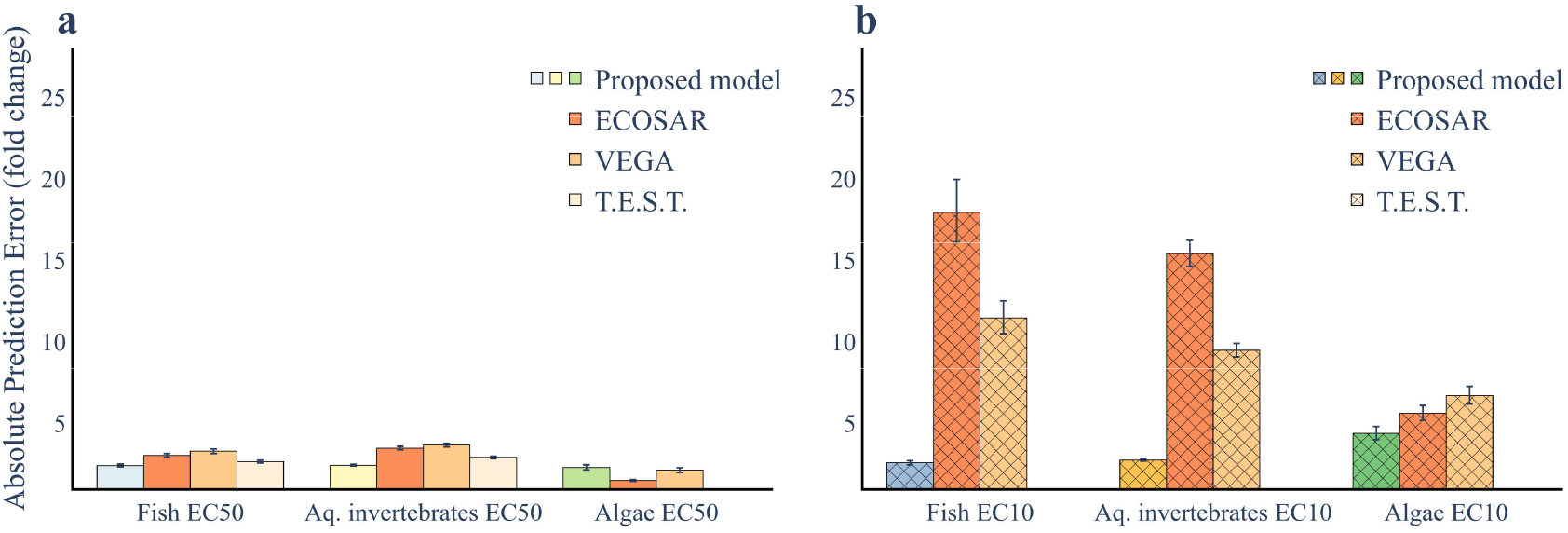
Comparison of performance of evaluated models. Panels (a,b) show the performance as the median absolute prediction error, measured as the absolute fold-change between predicted and experimental values, determined from the ten times repeated ten-fold cross-validations for the transformer-based model and the three QSAR methods, ECOSAR, VEGA, and T.E.S.T. The comparison was done only for chemicals within the applicability domains of all methods but not in the training datasets of any of the methods. In panel (a) models for EC_50_ (n are 297, 707, and 63 for fish, aquatic invertebrates, and algae, respectively). In panel (b) models for EC_10_ (n are 171, 598, and 116 for fish, aquatic invertebrates, and algae, respectively).

**Table 2:** Overview of model performance. Comparison of the model performance for the transformer-based model compared to ECOSAR, VEGA, and T.E.S.T. The number of data points in the intersect is the number of chemicals that are within the applicability domain of all models, but no model has been trained upon. The mean and median absolute error is calculated for this subset of chemicals for comparative purposes. The number of data points is the number of chemicals inside the applicability domain of each individual model, including training data for the QSAR methods but excluding training data for the transformer-based model. The three rightmost columns report the percentage of chemicals with an error larger than the respective fold difference.

| Dataset | Model | #Datapoints in the intersect | Mean absolute error | Median absolute error | #Datapoints | >1000 | >100 | >10 |
| --- | --- | --- | --- | --- | --- | --- | --- | --- |
| Fish EC<sub>50</sub> | Proposed Model | 297 | **3.37** | **2.40** | 1964 | **0.4** | **3.1** | **20.8** |
|  | ECOSAR | 297 | 4.20 | 3.01 | 927 | 3.9 | 9.5 | 25.8 |
|  | VEGA | 297 | 4.63 | 3.27 | 884 | 0.6 | 4.3 | 22.6 |
|  | T.E.S.T. | 297 | 4.09 | 2.64 | 654 | 2.8 | 10.7 | 29.5 |
| Fish EC<sub>10</sub> | Proposed Model | 171 | **3.74** | **2.56** | 2951 | **1.2** | **5.8** | **24.1** |
|  | ECOSAR | 171 | 23.51 | 17.94 | 1798 | 10.3 | 27.8 | 65.1 |
|  | VEGA | 171 | 14.84 | 11.46 | 197 | 4.1 | 12.7 | 51.8 |
|  | TEST | - | - | - | 0 | - | - | - |
| Aquatic invertebrates EC<sub>50</sub> | Proposed Model | 707 | **3.36** | **2.41** | 2893 | **0.8** | **3.7** | **20.6** |
|  | ECOSAR | 707 | 5.05 | 3.46 | 1674 | 4.5 | 10.0 | 31.1 |
|  | VEGA | 707 | 5.17 | 3.65 | 1194 | 0.9 | 5.9 | 27.6 |
|  | T.E.S.T. | 707 | 3.94 | 2.89 | 1477 | 1.1 | 5.3 | 25.3 |
| Aquatic invertebrates EC<sub>10</sub> | Proposed Model | 598 | **3.94** | **2.73** | 2913 | **1.1** | **6.5** | **25.4** |
|  | ECOSAR | 598 | 18.63 | 15.40 | 1818 | 6.1 | 21.1 | 56.3 |
|  | VEGA | 598 | 10.24 | 9.47 | 696 | 1.4 | 7.2 | 49.3 |
|  | T.E.S.T. | - | - | - | 0 | - | - | - |
| Algae EC<sub>50</sub> | Proposed Model | 63 | 3.83 | 2.27 | 2229 | 0.9 | 4.8 | **24.6** |
|  | ECOSAR | 63 | **1.90** | **1.47** | 475 | 2.7 | 7.6 | **24.6** |
|  | VEGA | 63 | 2.48 | 2.10 | 605 | **0.5** | **4.1** | 26.8 |
|  | T.E.S.T. | - | - | - | 0 | - | - | - |
| Algae EC<sub>10</sub> | Proposed Model | 116 | **6.15** | **4.37** | 2695 | 1.5 | 7.7 | **30.4** |
|  | ECOSAR | 116 | 7.84 | 5.60 | 810 | 3.7 | 11.7 | 36.3 |
|  | VEGA | 116 | 8.31 | 6.69 | 427 | **0.2** | **5.9** | 32.1 |
|  | T.E.S.T. | - | - | - | 0 | - | - | - |

Finally, the performance of all methods was assessed by investigating the residuals. In this analysis, we included all chemicals that were inside the applicability domain of each respective method, even those included in the training sets of the traditional QSAR methods. The results showed that the transformer-based model had the overall highest performance. For the prediction of EC_50_ for fish, only 8 (out of 1964, 0.4%) of the chemicals had an error larger than a factor 1000 and only 60 (3%) had an error larger than a factor 100 (Figure 7a). The corresponding error rates for the traditional QSAR methods ranged between 0.6%-3.9% and 4.3%-10.7% for a deviation of a factor of 1000 and 100, respectively (Figure 7b-d). For predictions of EC_10_ for fish, the differences were even more pronounced, and deviations of a factor of 1000 were between 3 to 9 times more likely for the traditional QSAR methods when compared to the transformer-based model (Figure 7e-h). Similar patterns could be seen for aquatic invertebrates and algae (S.I. Figure 6, S.I. Figure 7, S.I. Figure 8, Table 2). Except for the prediction of EC_50_ for algae the transformer-based model outperformed the other methods.

**Figure 7:**
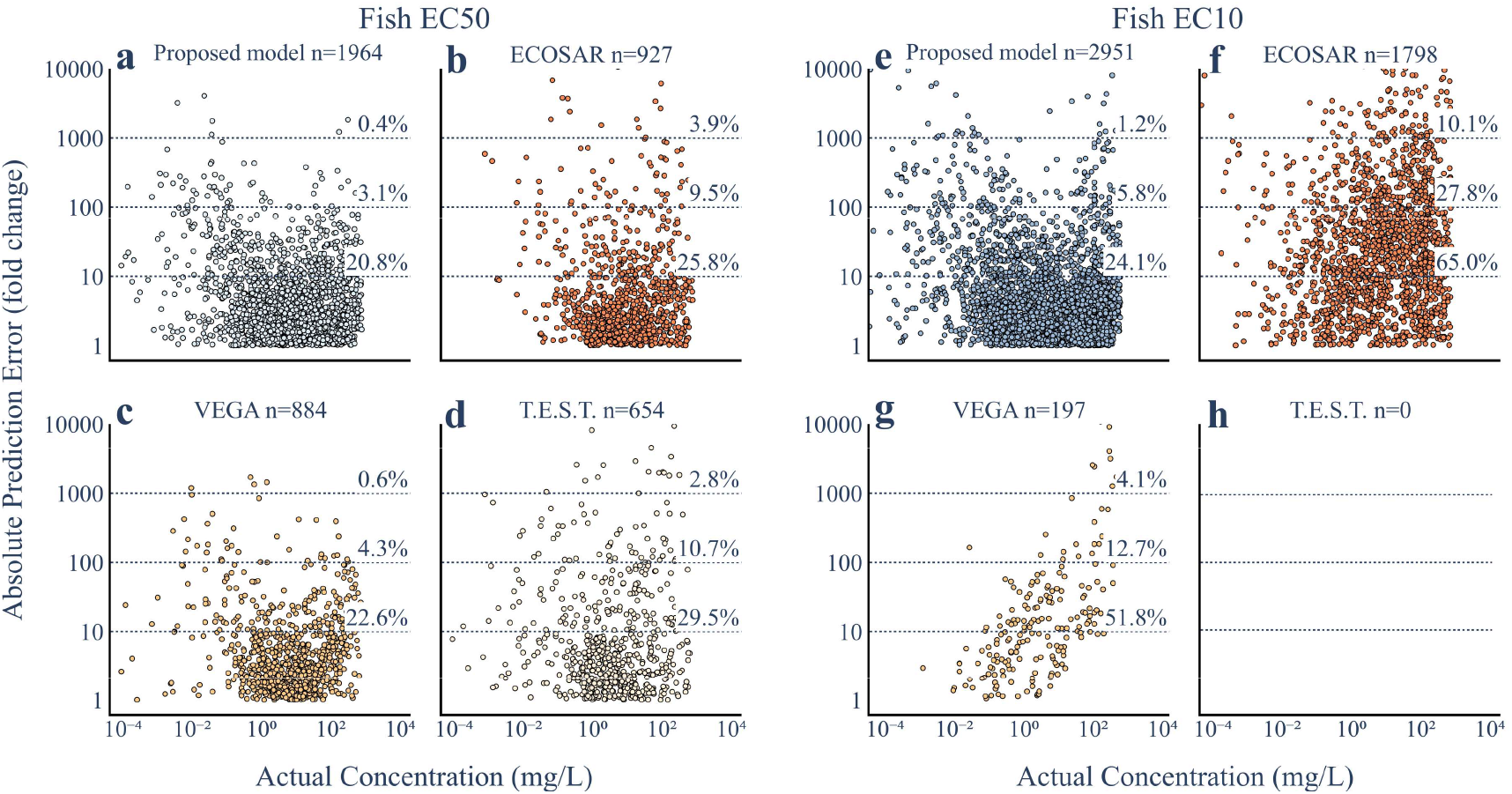
Absolute error distribution. The absolute prediction error against the experimental effect concentration for fish for all chemicals within the respective applicability domain of the transformer-based model, ECOSAR, VEGA, and T.E.S.T. Panels (a-d) show results for EC_50_ predictions and panels (e-h) show results for EC_10_ predictions.

## 3 Discussion

In this study, we show that deep learning techniques can significantly improve the computational predictions of chemical toxicity to aquatic organisms. We propose a model architecture that uses a transformer together with a deep neural network to derive the structure-toxicity and effect relationship directly from data. The resulting model has a large applicability domain and can make predictions for a diverse set of chemical classes. In addition, the proposed model has superior performance, with a median error corresponding to a factor of 2.00-4.11, when compared to experimentally measured effect concentrations. The high performance compared to traditional QSARs was especially pronounced for predictions of EC_10_, where the median error was up to a factor of 6.2 times lower. In addition to the benchmarking against ECOSAR, VEGA, and T.E.S.T. we also evaluated our model against another machine-learning model published in 2019, that only included fish (Sheffield and Judson, 2019). However, as that model is not publicly available, we here benchmarked by training our model using the same underlying data and the subsequent evaluation showed that our model performed on par for the reported measures of root mean square error and percentage of errors of different magnitudes.

One major difference between the proposed transformer-based model and more traditional QSAR methods is the numerical representation of the chemical structures. Our model uses self-attention, a mechanism originally used to infer complex dependencies in natural language data (Devlin et al., 2018; Vaswani et al., 2017), that has recently been shown to efficiently associate biochemical properties with molecular structure (Jumper et al., 2021). During the model training, the representation of the chemical structure is derived directly from the data by the transformer that up-weights the structural features that are especially important for toxicity. This can be contrasted with traditional QSAR methods. For example, for fish, ECOSAR v2.2 contains 263 models covering 111 chemical classes (Wright et al., 2022b). In addition, QSAR methods typically use pre-defined static structural representations, either using single values (e.g., hydrophobicity (Wright et al., 2022a)) or multi-dimensional vectors (e.g., fingerprints, or other vector molecular descriptors (Mayr et al., 2016; Sosnin et al., 2019)). However, our results demonstrate that self-attention constitutes an adaptive data-driven approach for estimating the structure-toxicity relationship, in turn improving the predictive performance beyond existing QSAR methods.

The model was trained on a dataset consisting of 143 829 experimental measurements for 6469 chemical structures in 1842 species. The high performance of the final model demonstrates that the currently available data is sufficiently rich to train deep learning models that are, at least on par, but often superior to existing methods in predicting chemical toxicity. This suggests that we have reached a tipping point within ecotoxicology, where data-driven black-box models are able to outcompete white-box QSAR approaches. In addition, this performance gap will likely continue to grow as more experimental data become available. We, therefore, argue that the adoption and further refinement of data-driven AI-based methods should be emphasized and that they have the potential to further improve computational performance in the field of (eco)toxicity. However, data-driven methods are dependent on the availability of large volumes of high-quality data. Therefore, a significant effort was spent aggregating data from several independent sources to construct a dataset of sufficient size. The final dataset includes results both from standardized tests and experiments performed by the scientific community. This effort mirrored the lack of standardized and organized ecotoxicity data and metadata which constitutes a major hurdle for the development and improvement of data-driven methods. Indeed, the data describing chemical toxicity is often both limited, incomplete, or inaccessible. For example, toxicity data generated to comply with European legislations, such as REACH, is only available as point estimates (typically EC_50_, EC_10,_ and no effect concentrations (NOECs)) while we see that extrapolation between effect concentrations is improved with increasing types of data. In addition, the data is currently not available in an easily accessible format. Meanwhile, results from toxicity experiments presented in scientific writing are not submitted to data repositories and are only manually extracted from the texts, making the data both hard to access and error-prone. Indeed, the US EPA estimates that they currently have a backlog of around 60,000 papers containing information that should be in the ECOTOX database (personal communication), one of the sources used in the model development. Additionally, several of the largest deviations between the measured and predicted concentrations were likely caused by manual data handling (see for example CAS 473-55-2, Pinane; (Passino and Smith, 1987)). Thus, adapting modern data-sharing strategies, based for example on the FAIR principle, in both academy, government, and industry would facilitate the development of more data-driven approaches within ecotoxicology.

Improved computational approaches will in turn help replace, reduce, and refine the use of animals for experimental purposes (Wilkinson et al., 2016). It will also allow rapid pre-screening of large and diverse bodies of data and aid in the development of more sustainable chemicals as well as facilitate the substitution to more benign ones (van Dijk et al., 2022). In turn, this will lower the societal cost, both from the testing of new chemicals, but also by reducing the burden of disease and impacts on ecosystem services from chemical pollution. Both issues that are becoming increasingly important as the number of chemicals in society grows and exposure to chemical mixtures becomes increasingly complex. In addition, only by employing computational methods will we ever be able to evaluate the toxicity of all the individual chemicals already in use (Wang et al., 2020) and it is also the only viable path for keeping up with the ever-expanding number of new ones.

## 4 Methods

### 4.1 Model description

The model consisted of two modules: a transformer encoder and a deep neural network (DNN). We used the pre-trained RoBERTa transformer (ChemBERTa), which consists of 6 encoders, 12 attention heads, and an embedding dimension of 768 (Chithrananda et al., 2020). Tokenization was done using a byte pair encoding (BPE) tokenizer. A beginning-of-string token, henceforth termed ‘CLS’ (classification) token, was added to all sequences during tokenization. All input sequences were then padded or truncated to a maximum length of 100 tokens. The input to the DNN consisted of the CLS token concatenated with the log_10_-transformed exposure duration (hours) and a binary vector indicating the type of toxicity endpoint and effect. The DNN consisted of multiple fully connected layers with a dropout probability of 0.2 and Rectified Linear Unit (ReLU) activation functions. The output layer consisted of a single node predicting the log_10_ effect concentration. The model was implemented in PyTorch v1.10.2 with ChemBERTa loaded from the Huggingface v4.21.1 transformers library.

### 4.2 Toxicity data

Data on the toxicity of chemicals towards fish, aquatic invertebrates, and algae was gathered from three sources, REACH dossiers, the US EPA database ECOTOX and the EFSA collection of pesticide registration data (EFSA, 2020; EPA, 2020; REACH Database, n.d.). Toxicity data in REACH was retrieved in August 2020 while the EFSA ‘openTox’ database and the US EPA ECOTOX database were retrieved in November 2020. The data was curated to harmonize concentration measures, annotations of limit tests, durations, effects, and endpoints and to remove all read-across data. Toxicity data from the three datasets were merged and all species names were verified using the R-package Taxize v0.9.99. The taxonomic groups for all species were harmonized to the same classification as used by the US EPA whenever possible. All no-effect concentration (NOEC) values and effect concentration (EC)/lethal concentration (LC) values reported between 0 and 10% effect were translated to EC_10_ values. All limit tests (data reported as ‘greater than’ or ‘less than’) were excluded from the analysis together with all reported effect concentrations larger than 500 mg/L. All training and validation were performed using log_10_-transformed test exposure and concentration data. All chemical structures, represented as SMILES, were collected by translating the reported CAS from the original datasets using the chemical identity resolver (R package webchem v1.1.2), collecting the first suggested SMILES. The SMILES were then canonicalized through RDKit v2022.03.5.

For algae, we only considered data from population toxicity assays (EC_50_ and EC_10_). For aquatic invertebrates, we considered assays that measured mortality (EC_50_, EC_10_), intoxication (EC_50_, EC_10_), reproduction (EC_10_), development (EC_10_), morphology (EC_10_), and population (EC_10_). For fish, the two datasets contained data for the same toxicity effects as the aquatic invertebrate dataset with the addition of growth (EC_10_).

### 4.3 Model Training

In total, nine individual models were trained. For each organism (algae, aquatic invertebrates, and fish), we trained models predicting EC_50_ and EC_10_ as well as a combined model predicting both EC_50_ and EC_10_. Model parameters (learning rate, batch size, number of reinitialized encoders, and number of hidden layers in the DNN) were set to be identical across organism groups. Thus, three parameter configurations were used in total (i.e., for prediction of EC_50_, EC_10_, and EC_50_/EC_10_, S.I Table 4). Specifically, all model configurations used a batch size of 512, three hidden layers (layer sizes 700, 500, and 300), and a learning rate set to 1.5E-4, 5.0E-4 or 2.0E-4 for prediction of EC_50_, EC_10_, and EC_50_/EC_10_ respectively. In total, the models contained between 84,490,145 and 84,495,745 trainable parameters. The used ChemBERTa transformer was pre-trained on 10 million SMILES as previously described (Chithrananda et al., 2020).

Model training was based on the mean absolute error (MAE) loss function using the AdamW (Loshchilov and Hutter, 2017) optimizer. The transformer encoder and the DNN were trained simultaneously. The learning rate was set to follow a linear schedule with a warmup phase, with a linear increase from 0 to the defined learning rate during 10% of the training steps and then linearly decreased. Layer-wise learning rate decay (LLRD) was employed based on the assumption that the first encoders capture very general language representations and that the last encoders are more task-specific (Zhang et al., 2020). During training, gradient norms were clipped to 1.0 to avoid exploding gradients (Zhang et al., 2019). All training was performed using an NVIDIA A100-SXM4-40GB GPU. Model hyperparameters were determined through Bayesian optimization (Weights and Biases v0.13.1.) based on five-fold cross-validation using the fish datasets (S.I. Figure 1, S.I. Table 1, S.I. Table 2, S.I. Table 3). Stratified sampling with probabilities inversely proportional to the number of data points for each combination of chemical structure, effect, and endpoint was used to account for skewness in the data.

### 4.4 Model Performance and Benchmarking

The models were trained using ten-fold cross-validation that was repeated ten times. For the models predicting EC_50_ and EC_10_, training and test datasets were split based on chemical structure. For the combined EC_50_/EC_10_, data was split based on pairs of effect concentration and chemical structure. The predictive performance was determined by first calculating the median residual for each unique combination of chemical, duration, effect, and endpoint. The overall model performance was then calculated as the weighted mean over all the combinations.

We compared our model against three commonly used QSAR-based methods; ECOSAR v2.2, VEGA v1.1.5, and T.E.S.T. v5.1.1.0 (Benfenati et al., 2013; Martin, 2020; Wright et al., 2022a). For the comparison with ECOSAR EC_50_ predictions, 96h fish (mortality), 48h invertebrate (mortality, intoxication), and 96h algae (population) measurements were used. For the comparison with VEGA EC_50_ predictions, 96h fish (mortality), 48h invertebrate (mortality, intoxication), and 72h algae (population) measurements were used. For ECOSAR the predicted ChV was also, in accordance with current guidelines, divided by the square root of 2 in order to estimate a predicted NOEC (ECHA, 2016). For the comparison with ECOSAR and VEGA NOEC predictions, EC_10_ measurements of all durations and effects were used. For the comparison with T.E.S.T., only 96h fish EC_50_ (mortality) and 48h invertebrate EC_50_ (mortality, intoxication) were used. For ECOSAR, when more than one value per chemical was reported for EC_50_ and ChV, the lowest reported respective value was collected (as suggested in the user manual (Wright et al., 2022a)). Furthermore, for VEGA when more than one value was reported, the prediction was taken as the value with the highest reliability score, i.e., ‘good’, ‘moderate’, and ‘low’, in that respective order.

The applicability domain (AD) of each method was evaluated based on two criteria. Firstly, counting all compounds where a prediction/experimental value was reported, even if the prediction was reported as outside of the AD. Secondly, only for compounds within the AD. For the performance comparison, the set of compounds inside the AD of all three QSARs, but excluding experimental/training data, was used (see S.I. ‘Supplementary Methods’ for details).

Finally, the model was compared to a machine learning-based model able to predict fish EC_50_ mortality and NOEC mortality and sub-lethal effects (Sheffield and Judson, 2019). However, as that model was not publicly available, our model was trained on the data included in Sheffield and Judson (2019) using ten-fold cross-validation and evaluated on the reported measures of root mean square error and percentage of errors of different magnitudes.

## Code availability

The code used to build, train, and evaluate the models presented in this study are available at https://github.com/StyrbjornKall/ecoCAIT.

## Acknowledgements

Funding from the FRAM Centre for Future Chemical Risk Assessment and Management at the University of Gothenburg was received by MG. SK. PS. JSI. JC. TB. SM. and EK. Funding from the Swedish Research Council was received by MG. JC and EK.

