## Supplementary Information for "Transformers enable accurate prediction of acute and chronic chemical toxicity in aquatic organisms"

### 1 Supplementary methods

#### 1.1 Determining QSAR applicability domains

T.E.S.T., VEGA and ECOSAR each have individual applicability domains and methods for reporting if a chemical is outside of that domain. T.E.S.T. only provides predictions for chemicals inside of the applicability domain. VEGA provides predictions for chemicals outside of the applicability domain but reports these as ‘low reliability’. ECOSAR reports with one or more ‘flags’ or ‘alerts’ if predictions are less reliable.

The QSAR dataset was constructed according to the following 1) for T.E.S.T., all reported predictions were included in the QSAR dataset. 2) for VEGA, all predictions with ‘low’ reliability were considered as outside of the applicability domain. 3) for ECOSAR all predictions with an associated ‘alert’ were considered as outside of the applicability domain, apart from when only the alerts, ‘AcuteToChronicRatios’ or ‘SaturateSolubility’ were reported as these were considered as less severe. Finally, for ECOSAR, predictions for chemicals reported as ‘SHOULD NOT BE PROFILED’ by ECOSAR v2.0 in batch mode were considered as outside of the applicability domain. (ECOSAR v2.0 was used as batch predictions from ECOSAR v2.2 report predictions without flags or alerts when predictions in single compound mode are reported as ‘SHOULD NOT BE PROFILED’). ECOSAR v2.0 was run using both CAS and SMILES as individual inputs to provide full coverage. This mostly reported metals and metal-containing salts.

#### 1.2 Extracting which chemicals the QSARs were trained with

Combinations of chemicals and species groups that the QSARs were trained on were labelled as follows. 1) For T.E.S.T. and VEGA all combinations with reported ‘experimental’ data were labelled. 2) For ECOSAR v2.2, the training data was retrieved from the file:

‘..\ecosarapplication v2_2\Helpful\Consolidated_SAR_MS.pdf’.

### 2 Supplementary results

#### 2.1 Selection of ChemBERTa version and loss function

Initially, we explored the available pre-trained versions of ChemBERTa and evaluated our model using two loss functions, mean absolute error (MAE/L1) and mean squared error (MSE). ChemBERTa is available with versions pre-trained using either a Byte-Pair Encoding (BPE) or a SMILES tokenizer and with varying amounts of SMILES included in the pre-training. The largest number of unique SMILES included in the pre-training was 10 and 1 million for the BPE and SMILES tokenizers, respectively. Hyperparameter sweeps using Bayesian optimization were performed for both versions, using either the MAE or MSE loss functions. Thus, the optimizer changed both the loss function and learning rate while minimizing the weighted median loss (S.I. Table 1). In total 30 individual models per ChemBERTa version were trained (i.e., testing ~15 different learning rates per version and loss function combination). Training was performed over 30 epochs, using five-fold cross-validation and the best-performing model was used for the evaluation (S.I. Figure 1). This optimization was performed using the largest individual dataset (fish $EC_{50}$).

**S.I. Table 1: ChemBERTa version and loss-function optimization.** Parameter/Hyperparameter sweep to determine the best performing ChemBERTa version and loss function based on the fish $EC_{50}$ dataset. This sweep uses a Bayesian optimizer and a five-fold cross-validation with the average median loss as the target metric. The choice of a log-uniform distribution for the learning rate ensures that learning rates from different magnitudes are equally likely to be tested.

<table>
<tr><th colspan="2"></th><th colspan="2">EC<sub>50</sub> model development</th></tr>
<tr><th>Parameter/Hyperparameter</th><th>Distribution</th><th>Interval</th><th>Result</th></tr>
<tr><td>Learning rate</td><td>Log-uniform</td><td>[1e-3,1e-6]</td><td>- [a]</td></tr>
<tr><td>ChemBERTa version</td><td>Categorical</td><td>[SMILES_tokenized_PubChem_shard00_160k, PubChem10M_SMILES_BPE_450k]</td><td>PubChem10M_SMILES_BPE_450k</td></tr>
<tr><td>Loss function</td><td>Categorical</td><td>[MAE Loss, MSE Loss]</td><td>MAE Loss</td></tr>
<tr><td colspan="4">FIXED VALUES</td></tr>
<tr><td colspan="2">Dataset</td><td colspan="2">Fish EC<sub>50</sub></td></tr>
<tr><td colspan="2">Epochs</td><td colspan="2">30</td></tr>
<tr><td colspan="2">Batch size</td><td colspan="2">512</td></tr>
<tr><td colspan="2">Dropout</td><td colspan="2">0.2</td></tr>
<tr><td colspan="2">Number of frozen encoders</td><td colspan="2">0</td></tr>
<tr><td colspan="2">Frozen embedding layer</td><td colspan="2">False</td></tr>
<tr><td colspan="2">Number of reinitialized encoders</td><td colspan="2">0</td></tr>
<tr><td colspan="2">Number of hidden layers</td><td colspan="2">2</td></tr>
<tr><td colspan="2">Hidden layer sizes</td><td colspan="2">[350,20]</td></tr>
</table>

[a] *The resulting learning rate is not specifically of interest, but an interval is necessary as the loss function varied between the tests.*

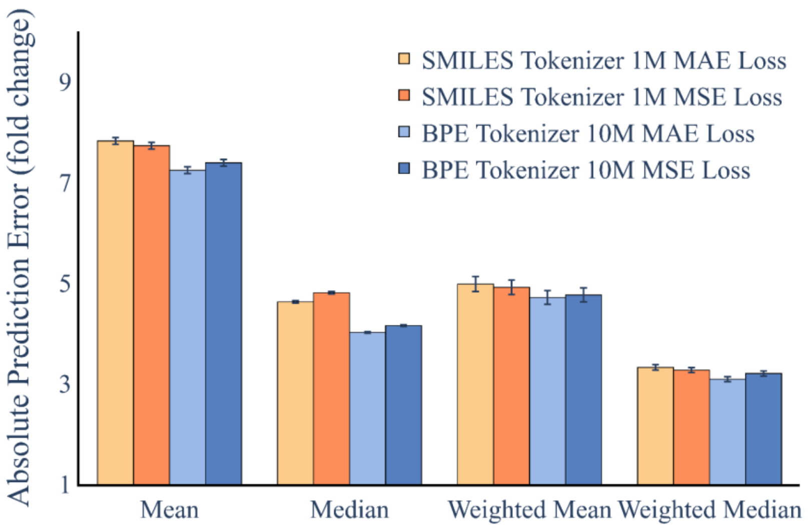

**S.I. Figure 1: ChemBERTa version and loss-function optimization.** Mean, median, weighted mean and weighted median validation loss for the five-fold cross-validation on the fish $EC_{50}$ dataset. Training and validation were performed using a transformer pre-trained either using a byte-pair (BPE) or SMILES tokenizer, and a pre-training-set of either 1 or 10 million SMILES. The error bars show the standard error of the mean when associated with mean errors, and the median absolute deviation (MAD) when associated with median errors. The validation losses were recorded at the epoch where the lowest normalized median validation loss was observed within each fold. The bars are based on the validations from the five 80/20 splits between training and validation. Thus, per fold n = 1934 SMILES was used for the training and n = 483 SMILES were used for the validation.

#### 2.2 Layer-wise Learning Rate Decay (LLRD)

Initial testing with the fish $EC_{50}$ and $EC_{10}$ datasets showed that layer-wise learning rate decay LLRD improved the fine-tuning stability and that LLRD had a very small influence on model performance. All models presented in the main paper are therefore trained using LLRD.

#### 2.3 Hyperparameter and parameter sweep setup

Initial testing showed that a fully trainable ChemBERTa transformer had the highest accuracy and consequently neither the embedding layer nor any of the encoders were frozen during the rest of the model development. Initial testing also showed that the dropout rate did not notably affect the accuracy and it was therefore fixed at 0.2.

The final model hyperparameter/-parameter configurations were determined by Bayesian optimization, training the model on the largest datasets per endpoint (fish $EC_{50}$ and $EC_{10}$). The optimization included the number of frozen and/or reinitialized encoders and frozen embedding layers in ChemBERTa, batch size, learning rate, dropout rate, as well as the number of hidden layers and number of neurons per layer for the deep neural network. Due to the high number of possible parameter combinations, the optimization was performed in two individual steps. The separation was performed based on the parameters either displaying a decoupling during initial training or because they had little or no impact on model accuracy during initial testing (S.I. Table 2, S.I. Table 3).

The resulting model configurations were decided based on overall performance (weighted median and mean error) and inter-model uniformity.

**S.I. Table 2: Batch size sweep.** Hyperparameter/parameter optimization to determine batch size based on the fish $EC_{50}$ dataset performed using Bayesian optimization with the average median loss across a five-fold cross-validation as the target metric. The choice of a log-uniform distribution for the learning rate ensures that learning rates from different magnitudes are equally likely to be tested.

| | | **$EC_{50}$ model development** | |
|---|---|---|---|
| **Parameter/Hyperparameter** | **Distribution** | **Interval** | **Result** |
| **Learning rate** | Log-uniform | [1e-3,1e-6] | - [a] |
| **Batch size** | Categorical | [512,256,128,64] | 512 |
| **FIXED VALUES** | | | |
| *Dataset* | *Fish $EC_{50}$* | | |
| *Epochs* | *30* | | |
| *Dropout* | *0.2* | | |
| *Number of frozen encoders* | *0* | | |
| *Frozen embedding layer* | *False* | | |
| *Number of reinitialized encoders* | *0* | | |
| *Number of hidden layers* | *2* | | |
| *Hidden layer sizes* | *[350,20]* | | |
| *ChemBERTa version* | *PubChem10M_SMILES_BPE_450k* | | |

[a] The resulting learning rate is not specifically of interest, but an interval is necessary as the batch sizes vary. The learning rate will therefore be subject to change in subsequent sweeps.

**S.I. Table 3: Learning rate, number of reinitialized encoders, hidden layers and hidden layer size optimization.** Hyperparameter/parameter optimization to determine the remaining parameter configurations for all three models. This is performed using Bayesian optimization with the average median loss across a five-fold cross-validation as the target metric. The choice of a log-uniform distribution for the learning rate ensures that learning rates from different magnitudes are equally likely to be tested.

| | | **$EC_{50}$ model development** | | **$EC_{10}$ model development** | | **$EC_{50}$/$EC_{10}$ combined model development** | |
|---|---|---|---|---|---|---|---|
| **Parameter/Hyperparameter** | **Distribution** | **Interval** | **Result** | **Interval** | **Result** | **Interval** | **Result** |
| **Learning rate** | Log-uniform | [1e-3,1e-6] | 1.5e-4 | [1e-3,1e-6] | 5e-4 | [1e-3,1e-6] | 2e-4 |
| **Number of reinitialized encoders** | Categorical | [0,1] | 0 | [0,1] | 0 | [0,1] | 0 |
| **Number of hidden layers** | Categorical | [2,3,4] | 3 | [2,3,4] | 3 | [2,3,4] | 3 |
| **First hidden layer size** | Int-uniform | [768,300] | 700 | [768,300] | 700 | [768,300] | 700 |
| **Subsequent hidden layer size** | Int-uniform | [500,20] | [500,300] | [500,20] | [500,300] | [500,20] | [500,300] |

**FIXED VALUES**

| | |
|---|---|
| *Dataset* | *Fish $EC_{50}$, Fish $EC_{10}$, and combined Fish $EC_{50}$ & Fish $EC_{10}$* |
| *Epochs* | *30* |
| *Batch size* | *512* |
| *Dropout* | *0.2* |
| *Number of frozen encoders* | *0* |
| *Frozen embedding layer* | *False* |
| *ChemBERTa version* | *PubChem10M_SMILES_BPE_450k* |

[a] The first hidden layer size is allowed a different interval than subsequent layers. Due to the definition of the sweep, subsequent hidden layers share the same interval but can assume different sizes. The result should be read as [size of sub. Layer 1, size of sub. Layer2]

#### 2.4 Hyperparameter and parameter sweep results

The final model parameter and hyperparameter configurations are presented in S.I. Table 4.

**S.I. Table 4: Final model hyperparameters.** The set of hyperparameters that yielded the highest performance for the model. The parameters were set after performing Bayesian optimization with Weights and Biases v0.13.1. The model configurations are used for all species groups and were derived from the largest individual dataset (fish $EC_{50}$, fish $EC_{10}$ and fish $EC_{50}/EC_{10}$).

| Model | Learning rate | Batch size | Number of reinitialized encoders | Number of hidden layers | Hidden layer sizes |
|---|---|---|---|---|---|
| **$EC_{50}$ model** | 1.5e-4 | 512 | 0 | 3 | [700, 500, 300] |
| **$EC_{10}$ model** | 5e-4 | 512 | 0 | 3 | [700, 500, 300] |
| **Combined $EC_{50}/EC_{10}$ model** | 2e-4 | 512 | 0 | 3 | [700, 500, 300] |

#### 2.5 Principal Component Analysis for invertebrates and algae

PCAs for the CLS-embeddings from the transformer when trained using the aquatic invertebrate $EC_{50}$ and $EC_{10}$ datasets (S.I. Figure 2) as well as the algae $EC_{50}$ and $EC_{10}$ datasets (S.I. Figure 3).

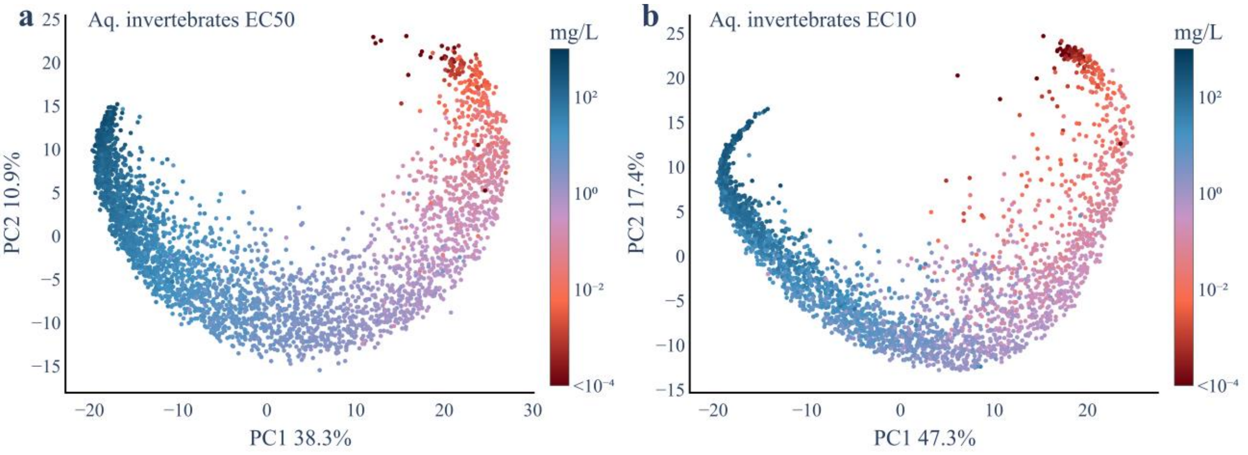

**S.I. Figure 2: PCA projection of CLS-embeddings from the transformer when trained on aquatic invertebrate $EC_{50}$ and $EC_{10}$ data.** Principal Component Analysis of CLS-embeddings from the transformer when trained using the (a) $EC_{50}$ dataset (b) $EC_{10}$ dataset.

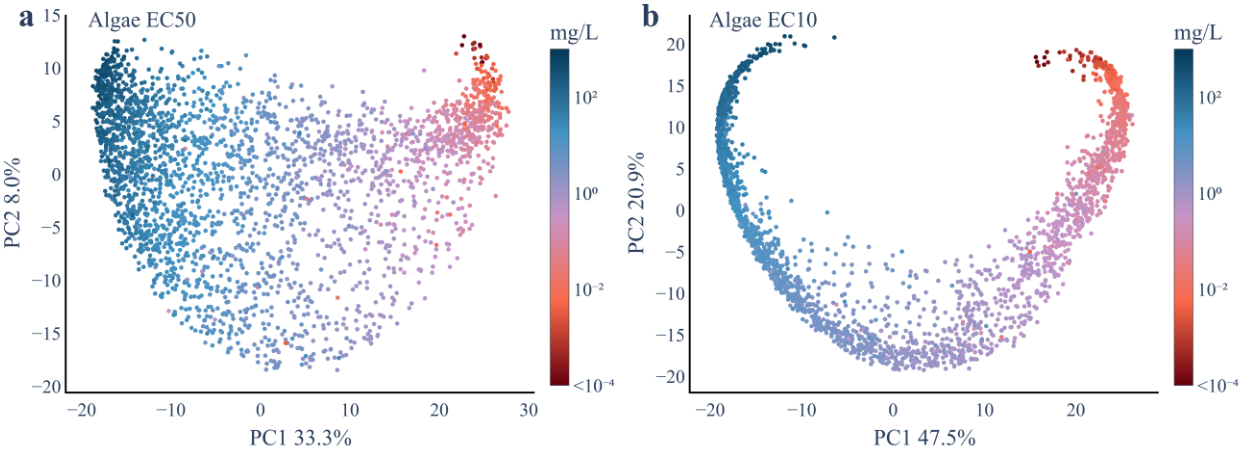

**S.I. Figure 3: PCA projection of CLS-embeddings from the transformer when trained on algae $EC_{50}$ and $EC_{10}$ data.** Principal Component Analysis of CLS-embeddings from the transformer when trained using the (a) $EC_{50}$ dataset (b) $EC_{10}$ dataset.

#### 2.6 Combined model comparison

The accuracy and residual distribution from the combined $EC_{50}$ and $EC_{10}$ model showed a consistent improvement across all species groups, demonstrating the model's ability to integrate and utilize different types of data (S.I. Figure 4). The improvement is seen both by the Absolute Prediction Error (fold change) and by a decrease in the number of residuals outside a factor of 10, 100 and 1000.

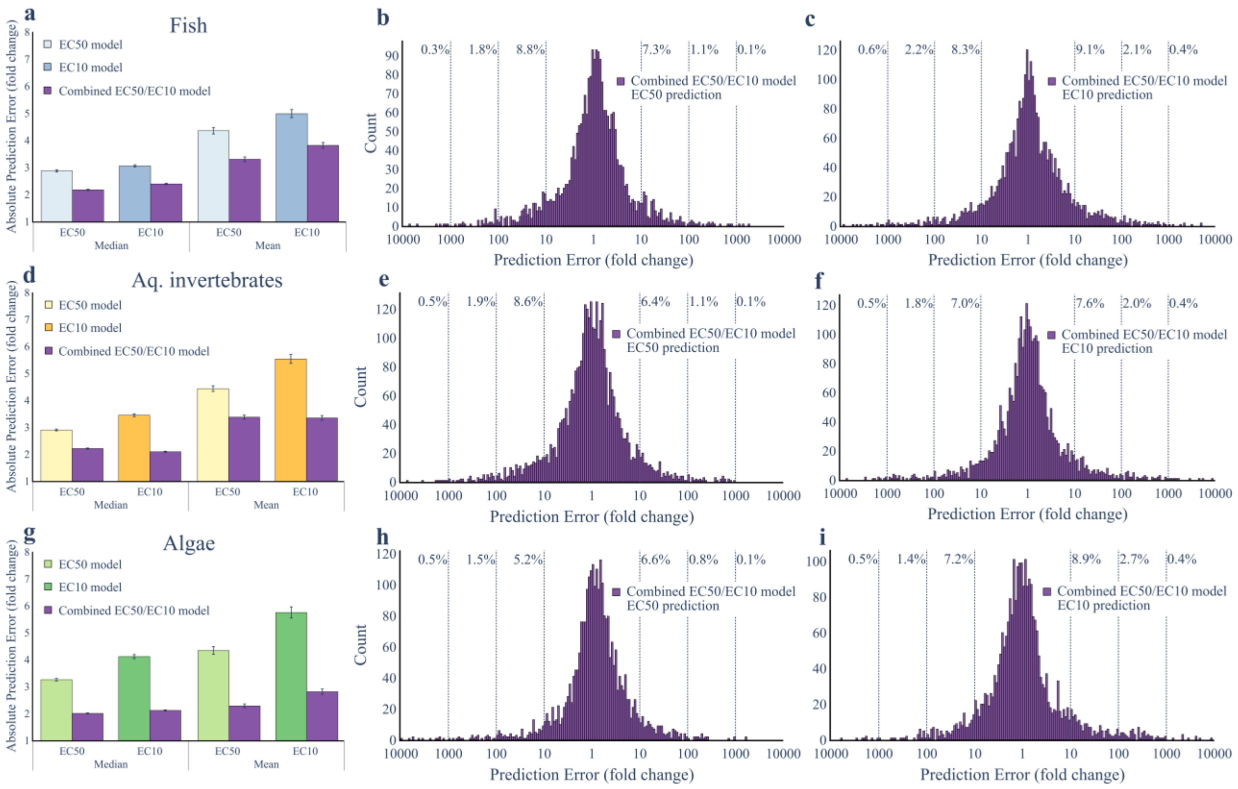

**S.I. Figure 4: Combined model performance fish, aquatic invertebrates, and algae.** Panels (a,d,g) show the performance as the absolute median and mean error, measured as the absolute fold-change between predicted and experimental values, determined from the ten-fold cross-validations for the (a) fish $EC_{50}$ model (n = 46104), $EC_{10}$ model (n = 23609), and the model able to predict both $EC_{50}/EC_{10}$ (n = 69713), (d) aquatic invertebrates $EC_{50}$ model (n = 31653), $EC_{10}$ model (n = 19170), and the model able to predict both $EC_{50}/EC_{10}$ (n = 54780), and (g) algae $EC_{50}$ model (n = 11909), $EC_{10}$ model (n = 11384), and the model able to predict both $EC_{50}/EC_{10}$ (n = 23293). The error bars show the median absolute deviation and the standard error of the mean for the respective prediction error. Panels (b-c, e-f, h-i) show the histogram of residuals for the (b-c) fish, (e-f) aquatic invertebrate, and (h-i) algae model able to predict both $EC_{50}/EC_{10}$, when predictions were evaluated on the $EC_{50}$ and $EC_{10}$ datasets. The reported percentage values show the percentage of chemicals which are erroneously predicted by a factor of more than 10, 100 or 1,000.

#### 2.7 Venn diagrams over QSAR applicability

The number of SMILES predictable by each QSAR method, for all chemicals with measured data, as well as all chemicals inside the QSARs applicability domains for the six individual datasets (S.I. Figure 5).

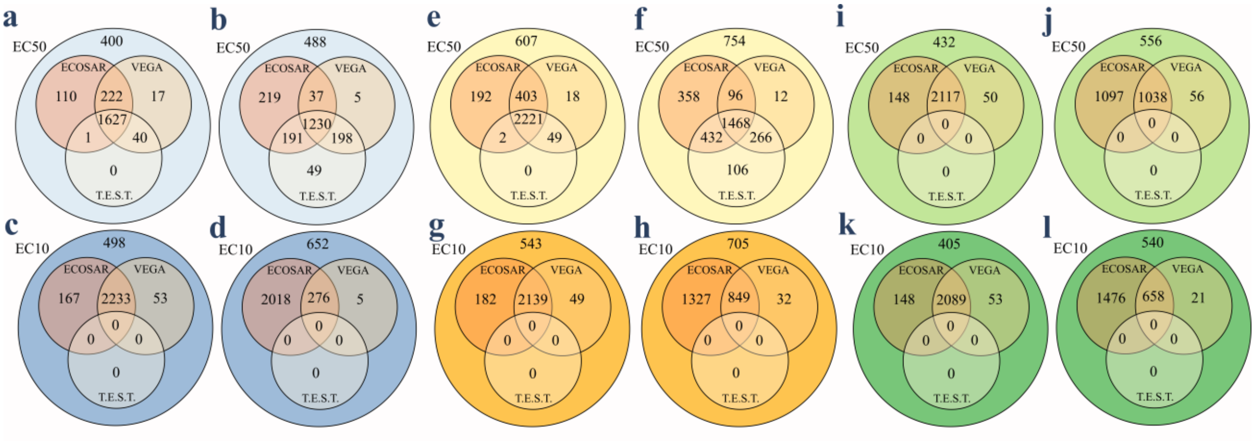

**S.I. Figure 5: QSAR models applicability intersection.** The number of SMILES predictable by each QSAR. (a,b) Number of predictable SMILES both in and outside of the applicability domain for the three QSARs based on all chemicals in the fish $EC_{50}$ dataset. (c,d) The number of predictable SMILES both in and outside of the applicability domain for the three QSARs based on all chemicals in the fish $EC_{10}$ dataset. (e,f) Number of predictable SMILES both in and outside of the applicability domain for the three QSAR tools based on all chemicals in the aquatic invertebrate $EC_{50}$ dataset. (g,h) Number of predictable SMILES both in and outside of the applicability domain for the three QSARs based on all chemicals in the aquatic invertebrate $EC_{10}$ dataset. (i,j) The number of predictable SMILES both in and outside of the applicability domain for the three QSARs based on all chemicals in the algae $EC_{50}$ dataset. (k,l) The number of predictable SMILES both in and outside of the applicability domain for the three QSARs based on all chemicals in the algae $EC_{10}$ dataset.

#### 2.8 QSAR accuracy and residual analysis

The accuracies were analyzed for the set of chemicals that were inside the shared applicability domain for all three QSARs (ECOSAR, VEGA, T.E.S.T.) for each of the six datasets, that neither model had been trained on. For our model the validation accuracy from the ten times repeated ten-fold cross-validation is used. Thus, neither model had been trained with the chemicals that are predicted, and no differences in coverage influence the comparison of accuracy. The residuals, per chemical, for all chemicals inside each model's applicability domain was also analyzed and results are summarized in the main text in Table 3. That figure is complemented here for fish (S.I. Figure 6), aquatic invertebrates (S.I. Figure 7) and algae (S.I. Figure 8).

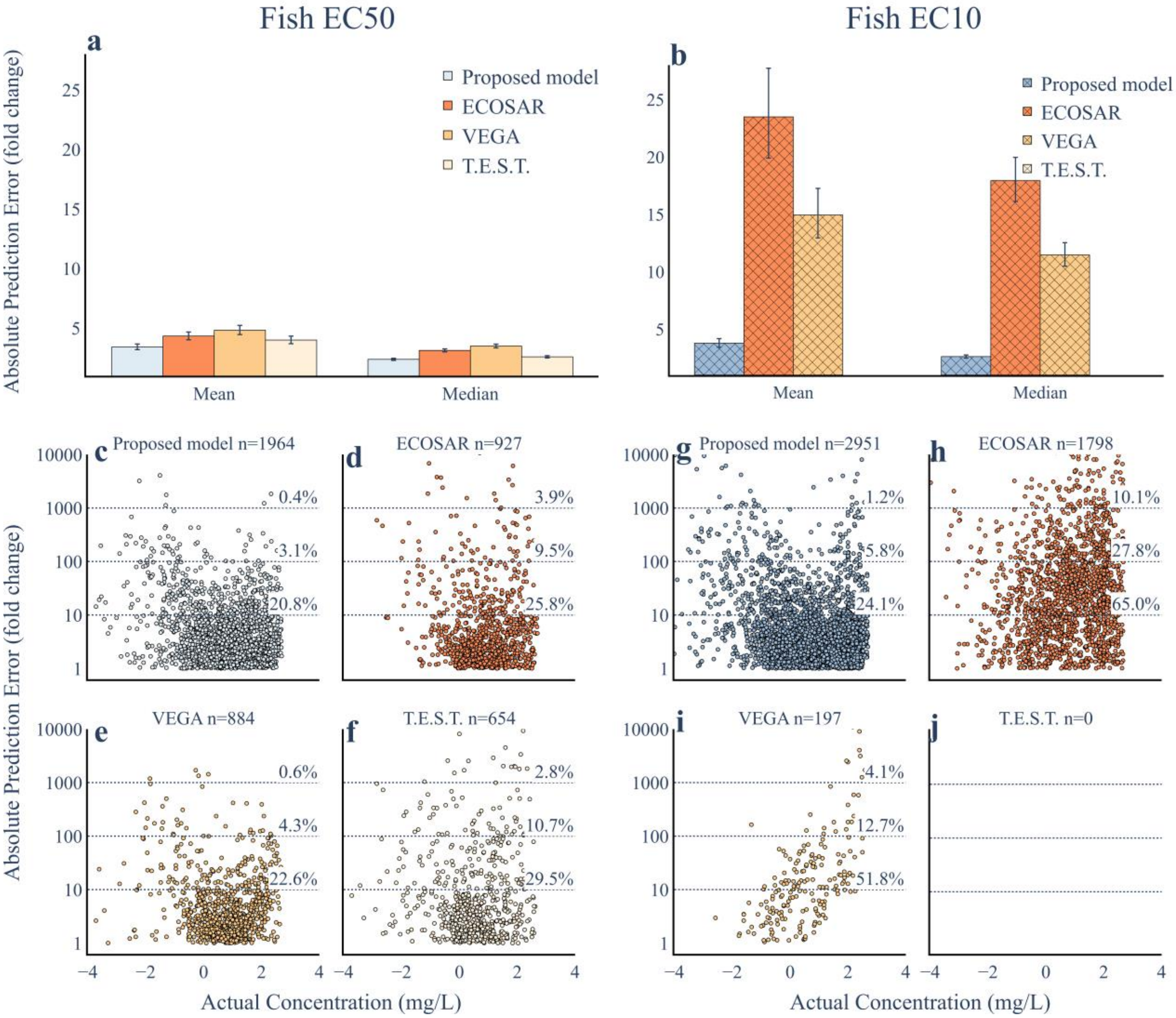

**S.I. Figure 6: Comparison of model performance and absolute error distribution fish.** The mean and median absolute error for the chemicals that are within the applicability domains, but not included in the training, of ECOSAR, VEGA and T.E.S.T. for the models trained using the (a) $EC_{50}$ dataset (n = 297) and (b) $EC_{10}$ dataset (n = 171). (c-j) The absolute error distribution for all chemicals within the applicability domain of the transformer-based model, ECOSAR, VEGA and T.E.S.T.

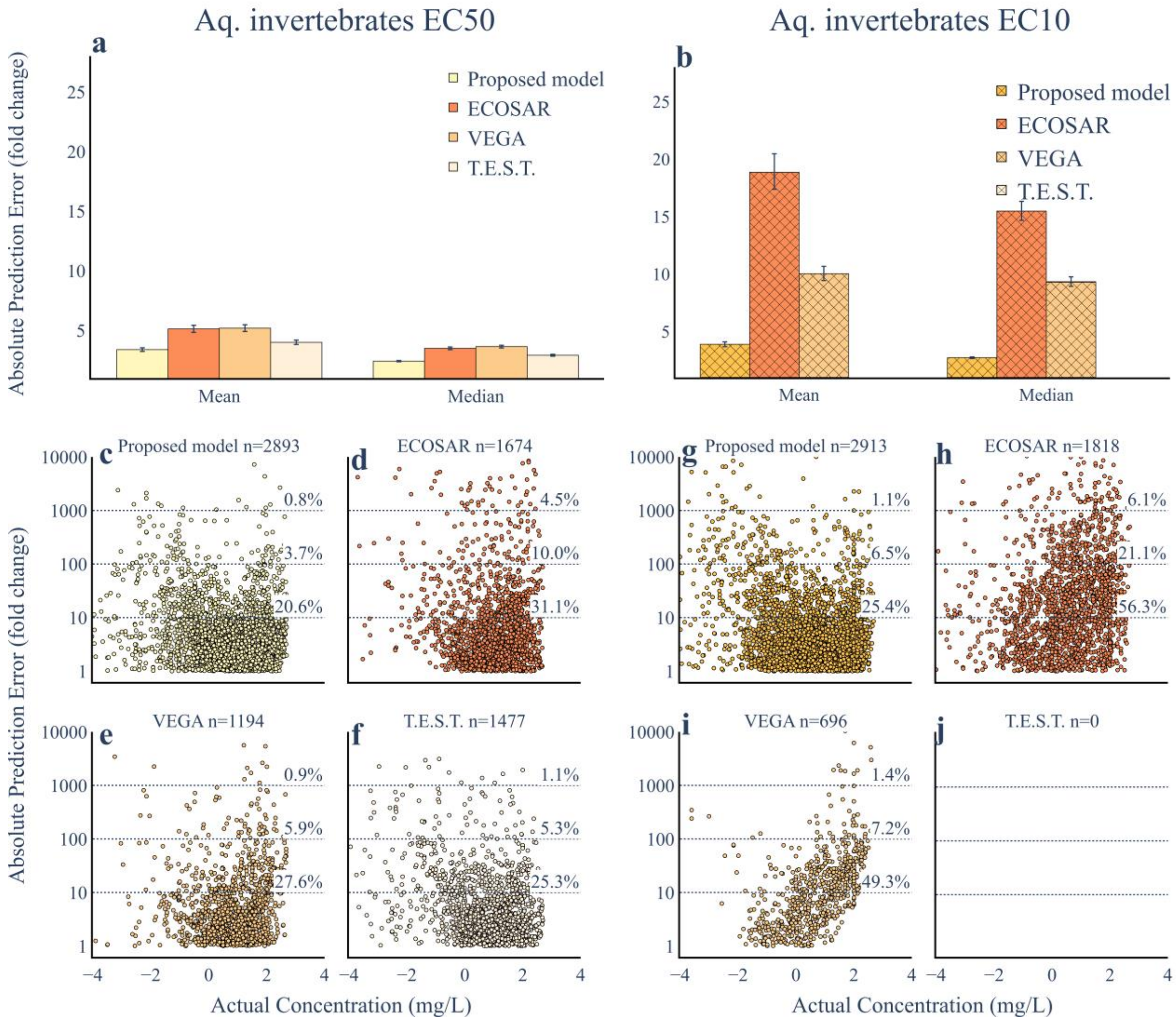

**S.I. Figure 7: Comparison of model performance and absolute error distribution aquatic invertebrates.** The mean and median absolute error for the chemicals that are within the applicability domains, but not included in the training, of ECOSAR, VEGA and T.E.S.T. for the models trained using the (a) $EC_{50}$ dataset (n = 707) and (b) $EC_{10}$ dataset (n = 598). (c-j) The absolute error distribution for all chemicals within the applicability domain of the transformer-based model, ECOSAR, VEGA and T.E.S.T.

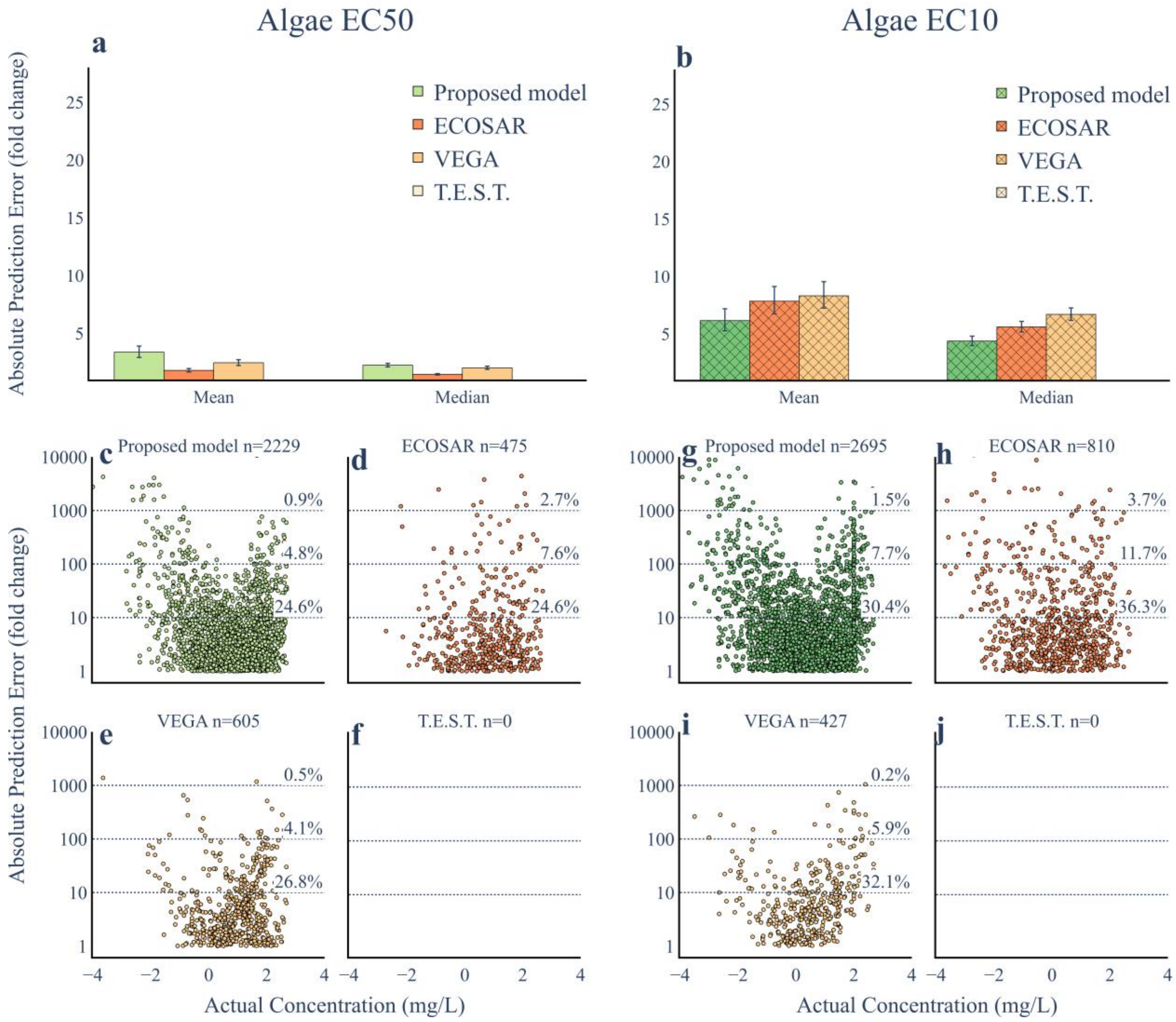

**S.I. Figure 8: Comparison of model performance and absolute error distribution algae.** The mean and median absolute error for the chemicals that are within the applicability domains, but not included in the training, of ECOSAR, VEGA and T.E.S.T. for the models trained using the (a) $EC_{50}$ dataset (n = 63) and (b) $EC_{10}$ dataset (n = 116). (c-j) The absolute error distribution for all chemicals within the applicability domain of the transformer-based model, ECOSAR, VEGA and T.E.S.T.

#### 2.9 QSAR residual analysis per effect for $EC_{10}$ datasets

The residual error was determined individually for each effect to ensure that the error distributions within the fish and aquatic invertebrate $EC_{10}$ datasets were not dependent on the measured effect (S.I. Figure 9, S.I. Figure 10). The proportions of errors exceeding 10, 100 and 1000 absolute fold prediction errors show that the model performs well also within each effect individually. Note, that algae are not presented as the model only predicts one (POP) effect.

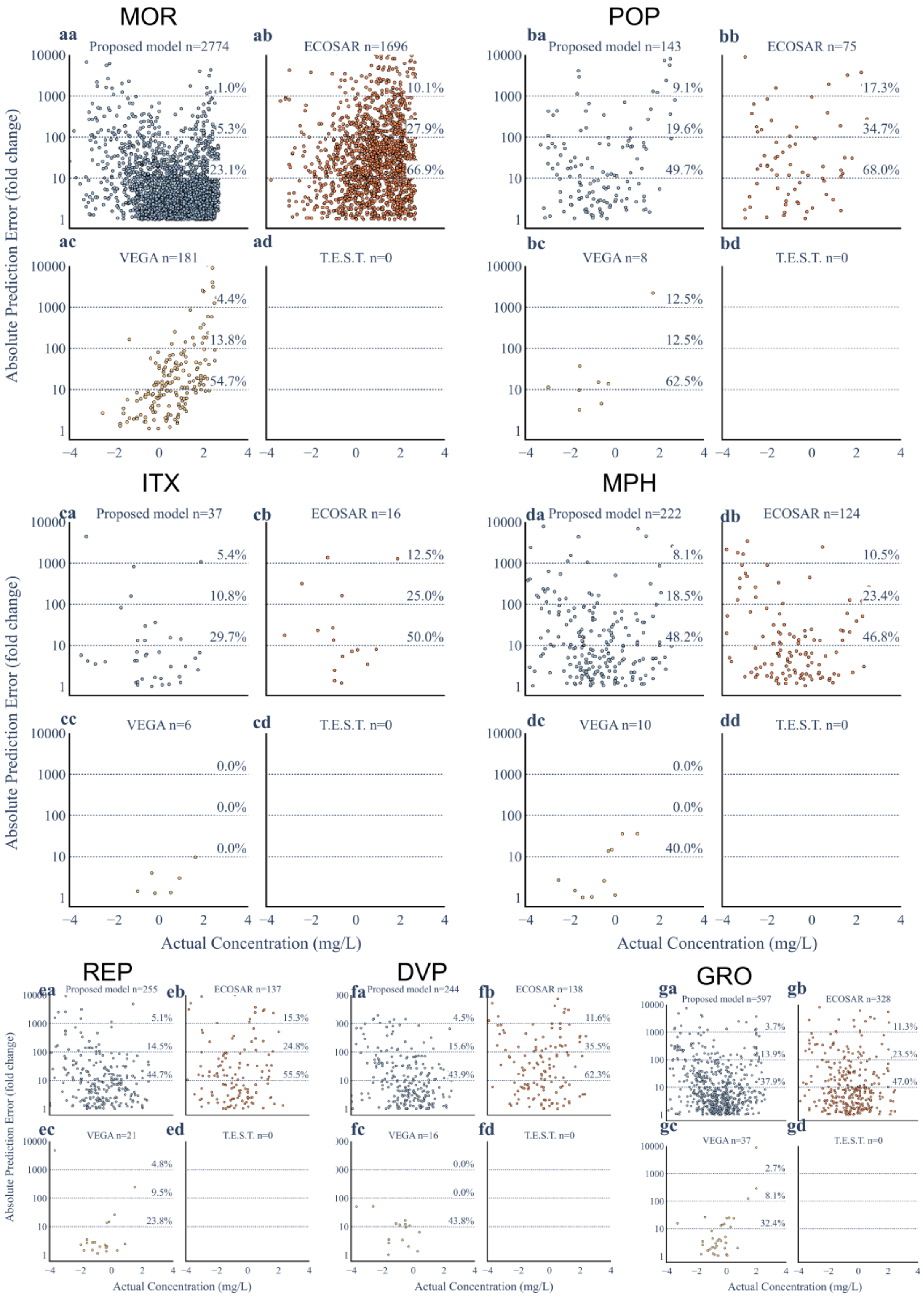

**S.I. Figure 9: Absolute error distribution per effect fish.** The absolute error distribution for all chemicals trained using the fish-$EC_{10}$ dataset for the transformer-based model, ECOSAR, VEGA and T.E.S.T., split by the effects that are inside the applicability domain of our model.

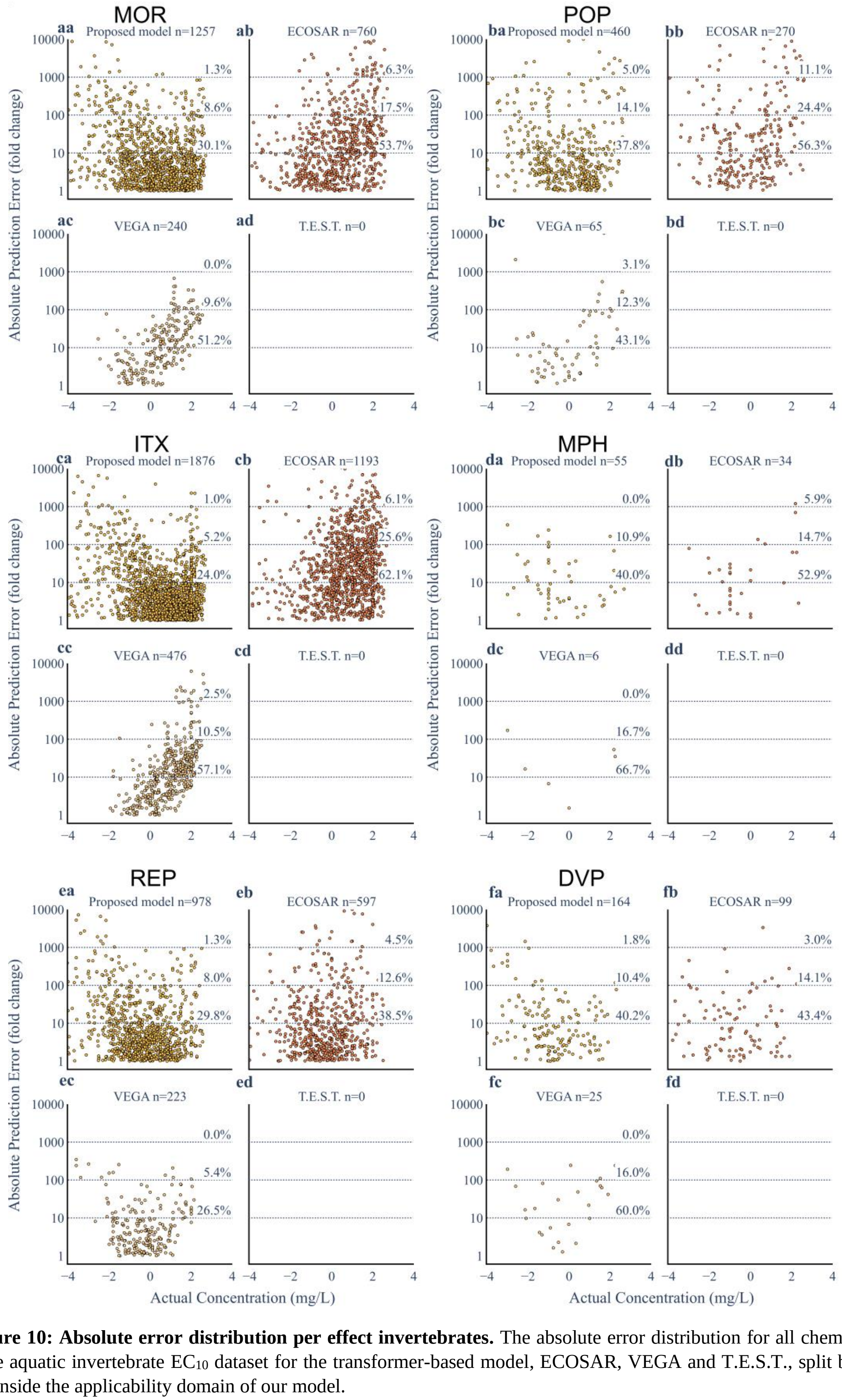

**S.I. Figure 10: Absolute error distribution per effect invertebrates.** The absolute error distribution for all chemicals trained using the aquatic invertebrate $EC_{10}$ dataset for the transformer-based model, ECOSAR, VEGA and T.E.S.T., split by the effects that are inside the applicability domain of our model.
